# Direct Synthesis of Targeted Nanosized ICG J-aggregate for Photoacoustic Imaging

**DOI:** 10.64898/2026.05.12.724349

**Authors:** Shrishti Singh, Leandro Soto Cordova, Nicholas Such, Marzieh Hanafi, Giovanni Giammanco, Dylan J. Lawrence, Iona E. Hill, Bijan Chamanara, Ilyas Fenaoui, Greeshma Tarimala, Dylan V. Scarton, Emna El Gazzah, Elsa Ronzier, Michael Girgis, Jeffrey L. Moran, Sunil Krishnan, Mariaelena Pierobon, Parag V. Chitnis, Remi Veneziano

## Abstract

Indocyanine green (ICG) J-aggregates (JAs) are self-assembled particles characterized by a sharp and strong absorption peak in the near-infrared region (∼890 nm), enhanced photostability, low fluorescence, and high photothermal conversion efficiency, compared to monomeric ICG. These attributes make ICG-JAs promising contrast agent candidates for photoacoustic imaging (PAI). However, traditional methods for synthesizing ICG-JAs often yield particles without targeting ability, which limit their applications. Thus, to synthesize targeted nanoscale JA, complex and multi-step encapsulation and filtration processes are generally required. To solve this issue, we introduce a robust and rapid strategy for direct synthesis of targeted nanoscale ICG-JA by co-assembling ICG and ICG-azide dyes under optimized formulation conditions that do not require encapsulation. The resulting nanoscale JAAZ particles (nJAAZ) exhibit diameters of ∼120-150 nm and are amenable to direct bio-orthogonal functionalization via copper-free click chemistry for the attachment of virtually any targeting ligands and/or biomolecules. We further demonstrate the strong photoacoustic signal generation of these nJAAZ in vitro and in vivo, highlighting their potential as a modular high-performance contrast agent platform for PAI. This work establishes a scalable and tunable platform for engineering functional JAs, opening new avenues for targeted molecular imaging and theranostic applications.

## Main

Photoacoustic imaging (PAI) is an emerging, non-invasive, hybrid imaging modality that combines optical excitation with acoustic detection^1–3^. PAI enables rapid acquisition of structural and functional information in biological tissues and can offer deeper tissue penetration along with higher sensitivity, contrast, and resolution compared to optical imaging alone^4–6^. In PAI, the signal is generated by chromophores embedded into tissues that undergo thermoelastic expansion upon absorbing pulsed laser light, subsequently emitting acoustic waves that are detected with ultrasound transducers^1,7^. These chromophores can be either endogenous biomolecules (e.g., hemoglobin and deoxyhemoglobin), which allow for label-free PAI, or exogenous contrast agents (e.g., gold nanospheres and organic fluorophores) delivered into the tissues to increase specificity, enhance contrast, and enable molecular imaging^6,8–12^. One of the most widely used exogenous contrast agents in PAI is the indocyanine green (ICG) dye, an FDA-approved cyanine dye already employed in several other preclinical and clinical fluorescence imaging and diagnostic applications^13–15^. ICG offers several advantages for PAI, including its strong near-infrared (NIR) absorption (peak monomeric excitation at ∼780 nm) and high photothermal conversion efficiency. These properties enable NIR-PAI for deep tissue imaging with reduced scattering and minimal absorption by endogenous chromophores^16–18^. However, ICG dyes suffer from limitations such as limited optical and structural stability, concentration dependent optical properties, and an inherent lack of targeting ability, which can drastically affect their imaging performance^19–21^.

Interestingly, like several other cyanine dyes, ICG can self-assemble into bricklike structures known as J-aggregates when incubated at elevated temperatures and high concentrations for several hours^22–26^. ICG J-aggregates (ICG-JA) provide superior photoacoustic performance over monomeric ICG through increased photostability, stronger and red–shifted NIR–I absorption (∼890 nm), low fluorescence, and enhanced photothermal conversion, enabling significantly deeper tissue imaging with PAI^27^. ICG JAs have been already successfully used for tumor imaging and image-guided therapy and have demonstrated superior imaging capability in term of depth of imaging and signal intensity in comparison with ICG^28–30^. However, since ICG-JAs are formed through a process of self-assembly, controlling their final size is challenging and their mean size often lies in the micron range with a broad size distribution, which can affect the breadth of their applications because of their limited capability to penetrate deep into tissue, to cross biological barrier and their reduced cellular uptake. Current methods to make targeted nanoscale ICG-JAs usually involve membrane filtration after synthesis to select the smaller particles^25^ and encapsulate a large amount of monomeric ICG in a targeted nanocarrier such as a liposome or a polymeric nanoparticle to create high local concentration leading to JA formation^29,31,28^. In general, these methods require multiple complex steps and are limited by the type of nanocarrier that can be utilized in the process. Moreover, adding a nanocarrier increases the complexity of ICG-JA synthesis, which can reduce its manufacturability and adds additional components that will be needed to meet clinical translation requirements for future clinical applications. Hence, developing a strategy for direct synthesis of nanosized ICG-JA with functional groups for further conjugation with targeting ligands would be more advantageous^32^.

In our previous studies^33,34^, we have introduced a method to synthesize modified ICG-JAs with sizes ranging from ∼0.25 to 1.2 µm. This approach also introduces azide (N₃) groups on their surfaces, enabling copper–free, click chemistry–based functionalization and we named these particles JAAZ^33^. In the present study, we modified our synthesis process to assemble sub-200 nm JAAZ particles, denoted as nJAAZ, ranging in size from 100 nm to 200 nm by optimizing key synthesis parameters including the molar ratio of ICG-N_3_:ICG dye and potassium chloride (KCl) concentration and without encapsulation or post-synthesis size selection (Figure 1). Cationic salts like KCl are known to influence the formation of J-aggregates of anionic cyanine dyes such as ICG^35,36^. Hence, in this study, we chose KCl and evaluated its importance in controlling the size of the J-aggregates in combination with other formulation parameters. Once synthesized and characterized, we demonstrate the potential of the nJAAZ platform for different imaging applications by conjugating various functional groups onto the azide groups at the surface of the particles. We notably show that the conjugation of a PEG layer can increase structural stability of the nJAAZ particles, the addition of folic acid can be used for targeting cancer cells, and the conjugation of fluorescent dyes such as Cy5 enables multimodal imaging and fluorescent-based tracking. Lastly, we evaluate concentration-dependent depth of imaging, biodistribution profile and PA signal intensity of the nJAAZ particles for in vivo imaging with a photoacoustic tomography instrument (TriTom® PhotoSound® Technologies, Inc., Houston, TX, USA). In conclusion, we have engineered a potentially translatable nanosized contrast agent platform that can be readily used in numerous PAI-based preclinical applications.

**Figure 1:**
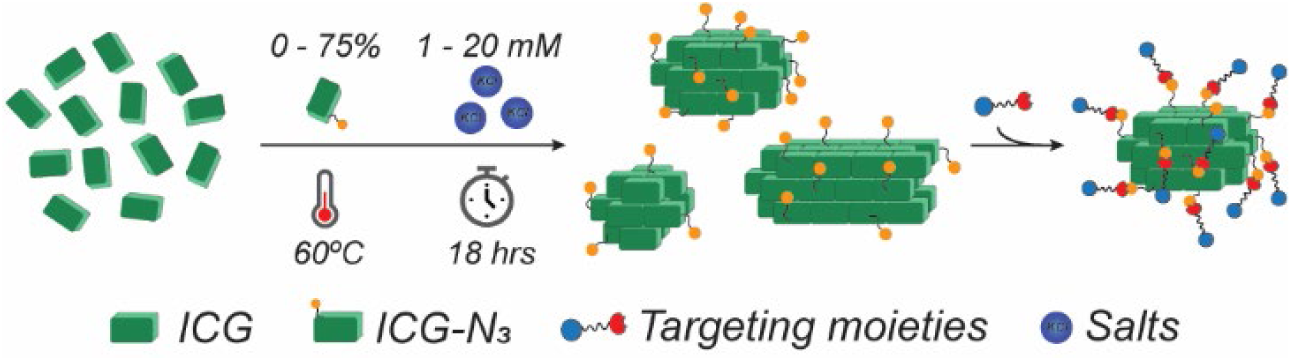
Targeted nJAAZ synthesis process. Graphical representation depicting the formulation and synthesis parameters controlled to tune the size of the J-aggregates composed of ICG and ICG-N_3_ dyes. A combination of increasing KCl concentration (from 1 to 20 mM) and increasing ratio of ICG-N_3_:ICG yields monodisperse, nanosized J-aggregates that can be directly modified via copper-free click chemistry with virtually any targeting moieties.

## Synthesis and characterization of the ICG-based nJAAZ particles

To synthetize the nJAAZ particles, we first evaluate the effect of two key synthesis parameters that were previously identified, namely the molar ratio of an azide modified ICG dye (ICG-N_3_) and ICG, and the KCl concentration^31^. Specifically, we are testing molar ratios of ICG-N_3_:ICG ranging from 1:10 to 1:1 at KCl concentrations ranging from 1 to 5 mM as our prior results indicate a direct relationship between KCl concentrations higher than 5 mM and the synthesis of submicron-sized ICG-JA particles. For the first set of experiments, the total dye concentration of ICG-N_3_ and ICG dye is kept at 250 µM for all ratios tested. To ensure completion of the aggregation reaction, we use a standard incubation protocol of 18 hours at a constant temperature of 60°C. To confirm the formation of J-aggregates we perform optical absorption measurements to detect the presence of the specific ICG-JA sharp absorbance peak at 895 nm^33^. For all ICG-N_3_:ICG ratios and KCl concentrations tested, we observe formation of JA with the characteristic absorbance peak at 895 nm (Figure 2a; Figure S1). The shift in absorbance is accompanied by a loss of fluorescence emission in the NIR region as observed with fluorescence spectroscopy using different excitation wavelengths (Figure S2). At lower KCl concentrations (1 and 2 mM), the 895 nm peak exhibits more pronounced intensity variations, with decreasing intensity correlating with increasing ICG–N₃ molar ratios. However, at 5 mM KCl, the intensity of the 895 nm peak remains largely unchanged across the different molar ratios, which is consistent with complete J-aggregate formation. Dynamic Light Scattering (DLS) is then used to determine the size of all ICG-JA formed after filtration using Amicon filters with a molecular weight cut-off of 150 kDa to remove any potential free dyes (Figure 2b; Figure S3; Table 1; Table S1). As previously shown, the hydrodynamic diameter of the JAs formed at a 1:10 molar ratio is ranging from 700 nm to ∼1100 nm for the highest concentration of KCl tested. When increasing the percentage of ICG-N_3_ (1:5 ICG-N_3_:ICG), the hydrodynamic diameter of the formed JAs decreased significantly with sizes ranging from 166 ± 19 nm using 1 mM KCl to 442 ± 125 nm when using 5 mM KCl (Figure S3). These results highlight the importance of both KCl concentrations and ratios of dyes for controlling the size of JAs. However, it is important to note that the JAs formed at a 1:5 molar ratio and all concentration of KCl tested appears highly polydisperse with a polydispersity index (PDI) superior to 0.5 (Table S1). Interestingly, when using a ratio of 1:1, we obtain sub-150 nm JA sizes with PDIs lower or equal to 0.2, for all KCl concentrations tested. The smaller size is obtained for 4 and 5 mM KCl with a 1:1 ratio of dyes with values of 121 and 120 nm, respectively (Figure 2b, Figure S3, and Table 1). These results are further confirmed with nanoparticle tracking analysis (NTA) measurements showing a highly monodispersed population of nJAAZ (1:1 ratio and 5 mM KCl) particles with an average size of ∼139.65 nm (Figure S4). With the NTA measurements we also determine the particles concentration estimated at 3.25×10^10^ particles/mL (for a total dye concentration determined at 250 µM). The difference between the average particle sizes measured by DLS and NTA can be explained by their respective weighting method, as DLS is intensity-weighted while NTA is number-weighted. Agarose gel electrophoresis (Fig 2b, inset) confirms the difference between the sizes of 1:10, 1:3, 2:5 and 1:1 ratio nJAAZ particles, with only JA particles formed at dye ratio lower than 1:3 able to enter in the dense gel and forming a sharp green band indicating monodisperse population.

**Figure 2:**
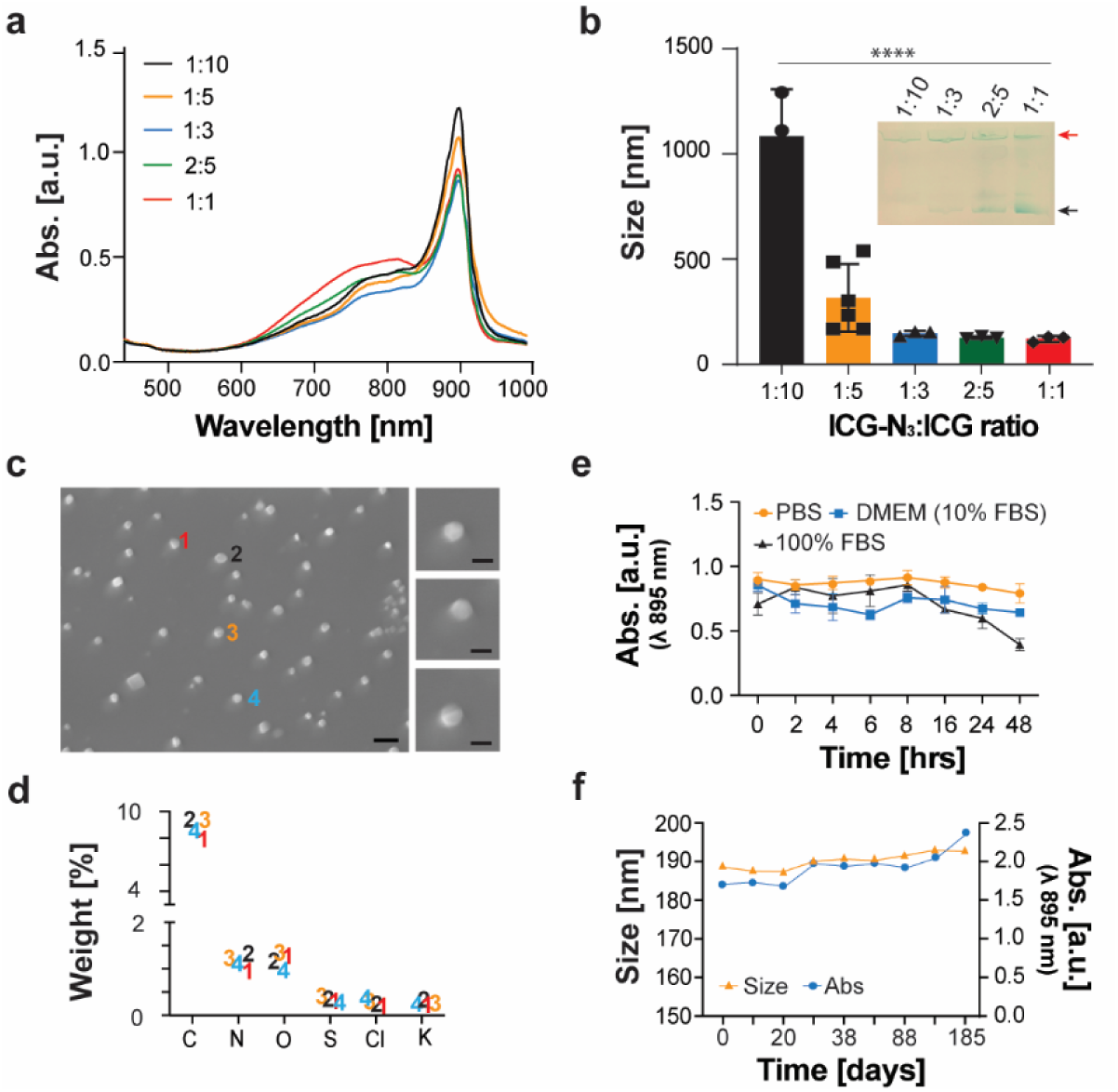
Characterization and stability of nJAAZ particles. (a) Absorbance spectrum of nJAAZ particles with varying ICG-N_3_:ICG molar ratios formed in 5 mM KCl. (b) Size of the formed nJAAZ particles and gel electrophoresis results of the samples with the red arrow indicating the gel wells and the black arrow the nJAAZ particles (inset). (c) SEM images and (d) EDS spectra of the nJAAZ particles. Scale bars are 250 nm (left panel) and 100 nm (right zoomed-in panels). (e) Stability of the nJAAZ particles in different buffers namely 1X PBS, DMEM+10% FBS and 100% FBS. (f) Absorbance and stability of the nJAAZ particles stored for 185 days in water. (n ≥ 3 individual replicates for a, b and e, p = **** = 0.0001).

**Table 1:** Hydrodynamic diameter measured by DLS of the different molar ratio ICG-N_3_:ICG samples.

| Molar ratios | KCl concentration [mM] |  |  |  |  |
| --- | --- | --- | --- | --- | --- |
|  | 1 | 2 | 3 | 4 | 5 |
| 1:10 | 735 ± 80 nm | 769 ± 33 nm | 795 ± 63 nm | 1089 ± 160 nm | 1085 ± 223 nm |
| 1:5 | 163 ± 19 nm | 176 ± 13 nm | 196 ± 5 nm | 304 ± 95 nm | 316 ± 161 nm |
| 1:3 | 161 ± 15 nm | 187 ± 9 nm | 181 ± 5 nm | 149 ± 21 nm | 142 ± 18 nm |
| 2:5 | 152 ± 17 nm | 137 ± 5 nm | 132 ± 10 nm | 132 ± 3 nm | 124 ± 18 nm |
| 1:1 | 146 ± 22 nm | 131 ± 15 nm | 126 ± 16 nm | 121 ± 17 nm | 120 ± 19 nm |

To further confirm incorporation of ICG-N_3_ in the JAs formed, Fourier transform infrared spectroscopy (FT-IR) is used with nJAAZ particles assembled at a molar ratio of 1:5 ICG-N_3_:ICG and 5 mM KCl. The graph presented in Figure S5 shows a peak at 2100 cm^-1^, which is typical of the azide group and similar to what we previously observed for the micron-sized JAAZ particles^33^. The percentage of incorporation of ICG azide is later confirmed by mass spectrometry with an incorporation rate of 50% for the nJAAZ assembled with the 1:1 ratio (Figure S6), which confirms that all dyes are properly incorporated into the aggregates. Scanning Electron Microscopy (SEM) images of nJAAZ particles reveal a spherical particle morphology with an average size of 104 ± 3 nm (Figure 2c) in line with our DLS and NTA measurements. Energy-dispersive spectroscopy (EDS) analysis (Figure 2d) showed the composition of the nJAAZ particles as carbon, nitrogen, oxygen and sulfur which match with the composition of ICG and ICG-N_3_ dyes. Altogether, these results indicate that an increase in the amount of ICG-N_3_ used in the formulation of the JAs associated with concentrations of KCl between 2 and 5 mM leads to the consistent formation of monodispersed nanosized J-aggregates. Moreover, our results confirm that tuning the ratios of dyes and the concentrations of KCl can be used to modulate the density of azide groups available per JA for subsequent surface modifications.

To better understand the process of formation of these aggregates, we also analyze the effect of the incubation time at a fixed concentration of KCl (5 mM) for different molar ratios of dyes. As seen in Figure S7a-c, the absorbance spectra show the typical JA peak at 895 nm as early as 6 hours for all dye molar ratios tested with size measured at 130 ± 15.9 nm, 122 ± 13.6 nm, and 125 ± 8.4 nm for 1:3, 2:5 and 1:1 molar ratios, respectively (Figure S7d-f). Moreover, the intensity of the 895 nm absorbance peak remains constant until 18 hours. These results mean that 6 hours is sufficient for the formation of the JAs, which could consequently simplify the manufacturing process.

The stability of nJAAZ particles is then assessed in vitro in phosphate buffered saline (PBS) at pH 7.4, in Dulbecco’s Modified Eagle Medium (DMEM) media supplemented with 10% fetal bovine serum (FBS), and in 100% FBS at 37°C. The nJAAZ particles are optically stable in PBS and in DMEM with 10% FBS for at least 48 hours with no decrease in the intensity of the absorbance peak at 895 nm (Figure 2e and Figure S8). DLS was not used to measure the size of the nJAAZ particles in serum and cell culture media due to the presence of many large biomolecules in these media that affect size readings. In the whole serum condition, a 50% decrease in the intensity of the absorbance peak at 895 nm is noted at 48 hours following a slow decrease of the intensity starting at around 8 hours (Figure 2e, black line and Figure S8b), which indicates a slow degradation of the nJAAZ particles. Overall, these results indicate that the nJAAZ particles retain their J-aggregate absorbance in PBS and serum-containing media over 48 hours, while partial signal loss in 100% serum suggests that serum-rich environments can affect long-term aggregate stability, yet they remain stable under physiological conditions for several hours and therefore could be used for in vivo imaging. The absorbance at 895 nm and size of nJAAZ particles are also recorded over 185 days to determine their long-term stability (Figure 2f). There is no significant decrease in the peak intensity or change in the size of the nJAAZs. Hence, our synthesized nJAAZ particles are stable when stored at 4 °C and protected from ambient light for up to 6 months without any changes in their physical properties.

## Functionalization of the nJAAZ particles

To enable targeting, the azide groups in the synthesized nJAAZ particles can be used for conjugation of various biomolecules or targeting moieties via copper-free click chemistry. In our previous work, we functionalized the JAAZ particles with streptavidin via biotin groups grafted onto the surface using a DBCO-biotin linker^33^. First, following a similar strategy, we first conjugate DBCO-biotin to the azide groups present on the surface of the nJAAZ particles and conjugate streptavidin after removing the excess of biotin groups via filtration. The conjugation efficiency is monitored by measuring the change in the ζ-potential of the nJAAZ particles. For the bare particles, the ζ-potential is measured at –56.6 ± 4.2 mV. The addition of DBCO-biotin increased the zeta potential value to –45.3 ± 5.4 mV. Moreover, the zeta potential changed to – 17.4 ± 2.4 mV when adding streptavidin (more neutral protein) to the surface (Figure S9a). In addition, the size of the bare, biotin and streptavidin functionalized nJAAZ did not significantly change (Figure S9b). The small but not significant increase in size following the addition of streptavidin could indicate efficient conjugation and confirm that the conjugation did not lead to aggregation of multiple JA particles.

In a second set of experiments, we further demonstrate the versatility of nJAAZ particles by directly functionalizing the azide groups with DBCO-nitriloacetic-acid (DBCO-NTA). The NTA group interacts with histidine (His) and can be used for the attachment of His-tagged proteins or peptides. Functionalization with DBCO-NTA caused a significant change in the value of the zeta potential of the samples at 1:10, 1:5, 1:3, 2:5 and 1:1 molar ratio that reaches values around -30 to -20 mV with the lowest value obtained for the 1:1 ratio (Figure S9c).

Finaly, we functionalized nJAAZ particles with folic acid, a folate receptor–targeting ligand for tumor-directed delivery. Surface azide groups on the particles are conjugated with DBCO–PEG–folate via copper-free click chemistry (Figure 3a). For all ratios tested, bare nJAAZ particles exhibit a strongly negative zeta potential (∼ –50 mV), which decrease in magnitude upon PEG–folate functionalization. Notably, the reduction in surface charge magnitude is more pronounced at higher ICG–N₃ ratios, consistent with increased PEG–folate conjugation resulting from a greater density of surface azide groups (Figure 3b). Particle size shows a small, non–significant increase following functionalization with DBCO–PEG–folate, as expected for the addition of a small ligand (Figure 3c). Surface plasmon resonance (SPR) confirms the targeting capability of folic acid–labeled nJAAZ (nJAAZ–FA) using a folate receptor (FOLR) immobilized on the sensor surface (Figure 3d). Binding to the receptor increased with nanoparticle concentration, indicating specific and concentration–dependent interactions.

**Figure 3:**
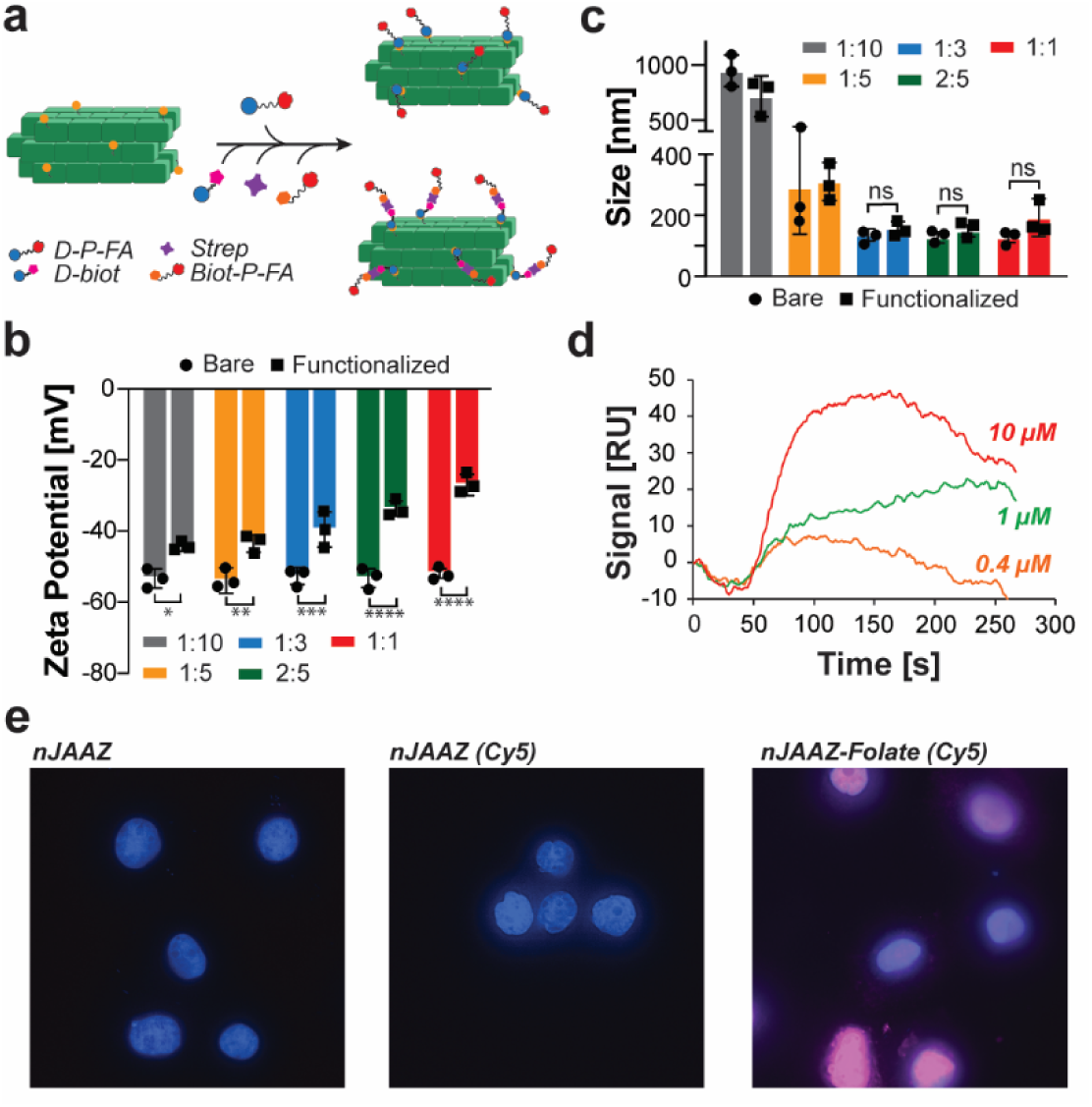
Functionalization of nJAAZ particles. (a) Schematic of copper-free click chemistry wherein compounds such as DBCO-PEG-folate (D-P-FA), DBCO-biotin (D-biot), streptavidin (Strep), and biotin_PEG-folic acid (Biot-P-FA) are conjugated to the surface of the nJAAZ particles onto the available azide groups (*in orange*) via direct conjugation or via streptavidin-biotin conjugation. (b) Zeta potential magnitude of different molar ratios of bare and DBCO-PEG-folate functionalized nJAAZ particles for different ICG-N_3_:ICG molar ratios. (c) Size of the different molar ratio JAAZ particles before and after functionalization with DBCO-PEG-folate. (n=3 for b and c (above), ns=non-significant, *= p<0.05, **= p<0.01, ***= p<0.001, **** p< 0.0001). (d). Representative SPR binding curves for the binding of nJAAZ-FA at 0.4, 1, and 10 µM on FOLR grafted on the SPR sensor. (e). Confocal fluorescence imaging of T47D cells incubated with various nJAAZ, fluorescently labelled nJAAZ, and fluorescently labelled nJAAZ-FA.

Targeting was further validated through cell–binding experiments using the FOLR–overexpressing breast cancer cell line model T47D previously used in FOLR targeted therapies^37,38^. To facilitate fluorescence imaging, nJAAZ particles are functionalized with a biotinylated folate ligand and visualized using Cy5–labeled streptavidin. As shown in Figure 3e, strong cellular binding is observed only for folic acid–functionalized nJAAZ particles, demonstrating receptor–specific association. Together, these results confirm the versatility of our platform and the effective folate–mediated targeting and demonstrate the feasibility of dual fluorescence and photoacoustic imaging using nJAAZ–FA as a multimodal contrast agent.

## Characterization of nJAAZ PA properties in vitro

We further evaluate the in vitro PA properties of the nJAAZ particles. The PA signal magnitude of the particles shows a dependence on the equivalent concentration of free dye, as seen with absorbance. With increased free dye concentration (Figure 4a), the PA signal proportionally increases for bare and folate functionalized nJAAZ particles. Similar results are obtained with the two other sizes tested (180 nm and 144 nm) and presented in Figure S10. The nJAAZ particles retain their 895 nm PA signal peak on the addition of folate molecules onto their surface (Figure 4b), identical to the bare particles. This signifies that addition of folate molecules does not alter the PA properties of the nJAAZ particles. For the further use of nJAAZ particles as a contrast agent, it is important to record the PA signal of the particles when mixed with whole blood. Even at low concentrations (∼12 µM) of equivalent free dye, the PA signal magnitude of both bare and functionalized nJAAZ particles is ∼2.5 times greater than the PA signal of whole blood (Figure 4c). Therefore, the nJAAZ can be used as a contrast agent since it can be detected against a common background chromophore such as hemoglobin. To evaluate the suitability of targeted nJAAZ particles for photoacoustic (PA) imaging, HeLa cells, which are also overexpressing the folate receptor^39^, were cultured on a glass slide patterned with an “M” shape to create a cell-free background reference in the shape of an “M” and cell growing around the “M” shape. As shown in Figure 4d, targeted nJAAZ particles exhibited strong binding to HeLa cells, resulting in a high PA signal in the cell region in comparison to the background. In contrast, non-targeted nJAAZ particles showed minimal signal, indicating negligible binding likely arising from nonspecific interactions.

**Figure 4:**
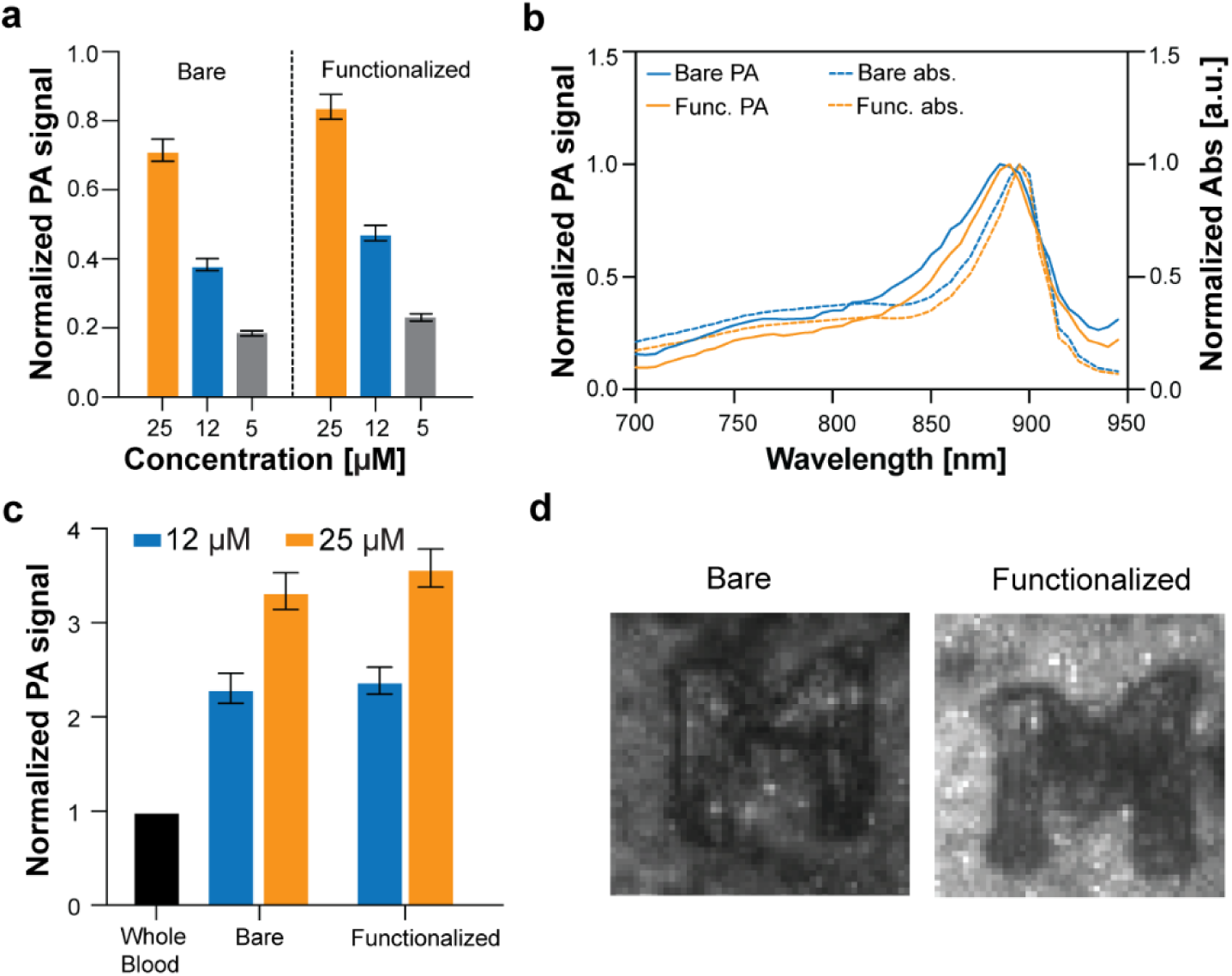
Photoacoustic properties of nJAAZ particles. (a) Photoacoustic signal magnitude of bare nJAAZ (size: 120 nm) and folate functionalized (nJAAZ-FA, size: 125 nm) particles at different equivalent concentrations of free dye. The data is normalized with respect to 1:100 ratio of India Ink used as a control. (b) PA spectrum *(solid lines)* and absorbance spectrum *(dashed lines)* of bare and functionalized nJAAZ particles. (c) PA signal magnitude of bare and functionalized nJAAZ particles mixed in whole blood. The signal is normalized with respect to whole blood. (d) PA images of nJAAZ *(left)* and nJAAZ-FA *(right)* targeted HeLa cells grown on a glass slide containing an “M” shaped structure to serve as background of the image were cells are not growing.

## In vivo PAI tomography imaging with nJAAZ

We further perform in vivo PAI using our nJAAZ contrast agent to demonstrate its potential for preclinical studies and future clinical translation. Multi-wavelength images were acquired at different timepoints using a commercial small animal photoacoustic imaging system, the TriTom® from PhotoSound, to understand the biodistribution, timescale of imaging, depth of imaging, and contrast-to-noise ratio of nJAAZ. Coronal maximum intensity projections of the mouse trunk are generated for three timepoints (0, 30, and 60 minutes) (Figure 5a and Figure S11). Interestingly, comparing the 150 µM and 400 µM volumes, greater photoacoustic signal retention is apparent for the lower concentration within the liver and spleen. One possible explanation is concentration-dependent uptake and clearance by liver and spleen macrophage populations in combination with signal masking from saturated superficial vasculature signal. Additional biodistribution studies would be required to test the macrophage clearance mechanism. To understand if superficial vasculature is masking deeper organ signal, the liver was segmented to exclude superficial vasculature for the 400 µM volume. At T60 minutes, the hepatocyte signal in the liver can be seen indicating that there is vasculature shielding present (Figure S12). This suggests that lower concentrations of nJAAZ may be ideal for in vivo imaging at time scales longer than half an hour. Next, 0.5 mm cross-sectional maximum intensity projections are generated at 3 different axial positions, T60 minutes post-injection to highlight anatomical features (Fig 5b). From the projection, superficial vasculature has the strongest PA signal as it is shallower from the skin line. In contrast, the thoracic vertebrae, liver, and spleen, are identifiable at a deeper depth in tissue. To confirm that the signal in the liver and spleen is due to nJAAZ, linear spectral unmixing is performed for oxyhemoglobin, deoxyhemoglobin, and nJAAZ (Figure 5c). The spectrally unmixed volumes at the initial time point show that nJAAZ is seen circulating through the blood stream of superficial vasculature. At T30 minutes post injection, nJAAZ can still be seen in vasculature while also beginning to accumulate in blood-rich organs such as the liver, spleen, and intestines, which is further supported by the results presented in Figure S12. At T60 minutes post injection, nJAAZ signal is primarily retained in the liver and spleen. This indicates that nJAAZ is contributing to PA signal deeper in the liver and spleen, and that PAI can be done 60 minutes post-injection. The iliac artery is used to calculate the contrast-to-noise ratio (CNR) and the respective imaging depth (Figure 5d). At an imaging depth of 5.2 mm, the CNR is found to be 8.0316. This is a three-fold improvement from (2.42 at 5 mm depth)^33^.

**Figure 5:**
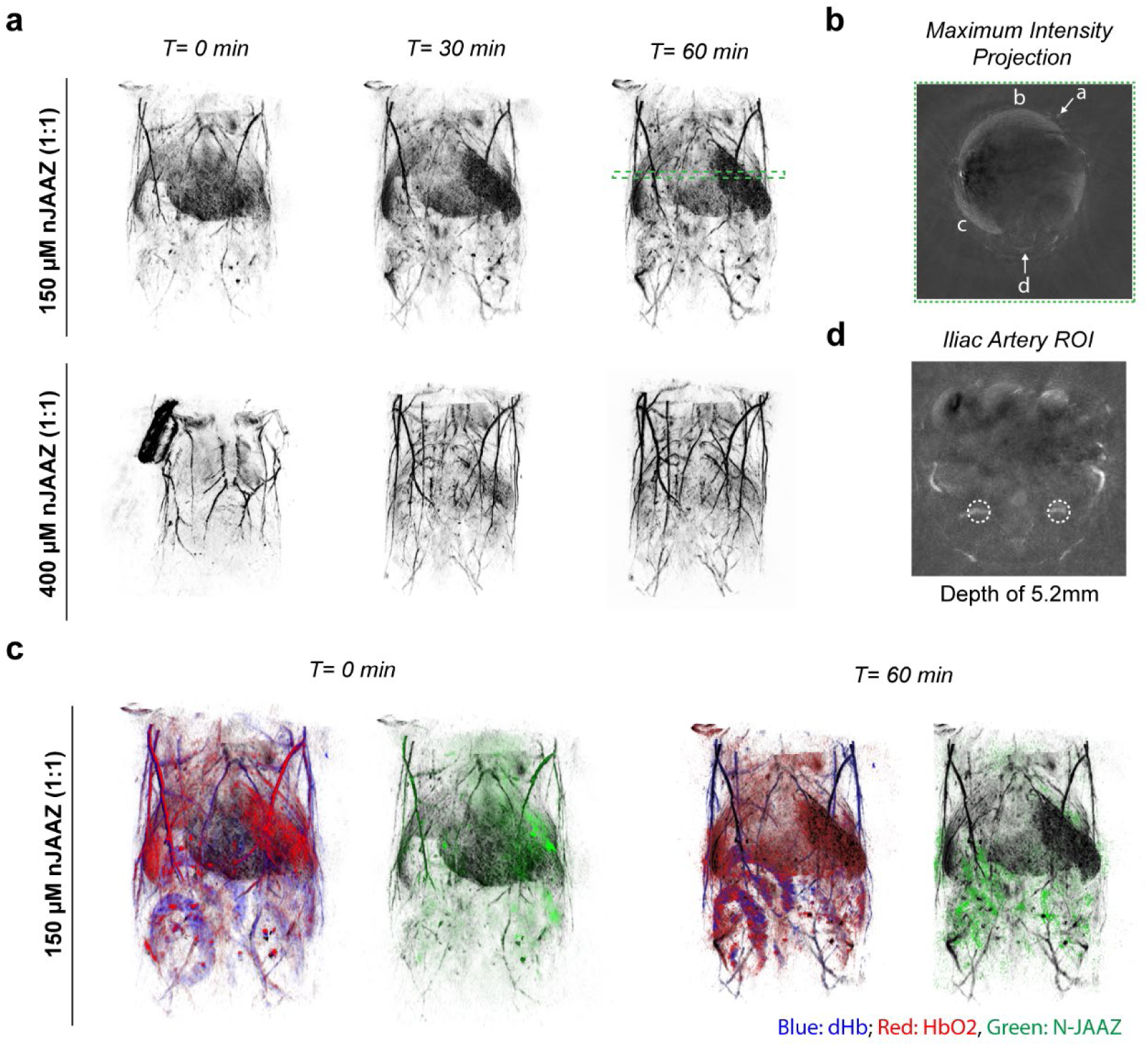
In vivo photoacoustic imaging of nJAAZ particles using a wavelength of 895nm. (a) Coronal slab maximum intensity projections of mouse trunk for 150 µM and 400 µM nJAAZ injections at three timepoints, post-injection: T0, T30, and T60 minutes. (**b**) Maximum intensity projection of an axial plane with a thickness of 0.5 mm. Highlighted features: a. superficial vasculature, b. liver, c. spleen, and d. thoracic vertebrae. In *ii*, the dashed circle highlights the iliac artery. (**c**) Coronal slab of spectrally unmixed deoxyhemoglobin (blue), oxyhemoglobin (red), and nJAAZ (green) at two timepoints: 0 and 60 minutes post-injection. Unmixed signals are overlayed over base 895 nm scan (black) to highlight anatomy. (d) Maximum intensity projection of an axial plane with a thickness of 0.5 mm with the iliac arteries shown with a dashed white circle.

## Conclusion and outlook

We demonstrate a robust, modular, and highly scalable synthesis method to create a nanoscale indocyanine green–based J–aggregates (nJAAZ) platform offering robust optical properties, size tunability, and versatile surface chemistry for efficient targeted NIR photoacoustic imaging. By systematically tuning the synthesis conditions and the relative ratio of ICG and ICG-N_3_ dyes, we can achieve highly monodisperse nanoparticles with controllable densities of surface–accessible azide groups. This level of synthetic control enables reproducible particle formation and efficient post–synthesis conjugation without compromising the hallmark J–aggregate optical signature in the NIR region or homogeneity in size distribution upon aggregation^31,32^. This is validated through various conjugation strategies, including direct copper-free click chemistry with DBCO conjugated folic acid and via streptavidin-biotin conjugation, which enabled receptor–specific binding in both surface plasmon resonance assays and tumor cell–based models, underscoring the feasibility of molecularly targeted nJAAZ constructs.

Additionally, a comparison experiment of the PA signal intensity between an nJAAZ (148 nm) and a micron-sized JAAZ sample (1120 nm) is performed to assess the size effect on the signal intensity. As presented in Figure S13, the nJAAZ particles provide about a two-fold higher PA signal magnitude regardless of the total dye concentrations used (15 and 50 µM) in comparison to their larger-sized counterpart at the same concentrations, which indicates that nJAAZ particles might provide stronger PA contrast in vivo. The higher photoacoustic signal from smaller aggregates could be attributed to more effective light absorption due to their higher surface-to-volume ratio, which increases dye exposure to incident light. These results indicate that the nJAAZ particles offer superior photoacoustic performance relative to previously reported micron–scale J–aggregates, yielding higher signal intensities at lower dye concentrations. This enhancement, combined with excellent signal against blood (2.5-fold higher than whole blood at 12 µM total dye concentration) and sustained stability under physiological conditions, enabled high–contrast in vivo photoacoustic tomography at more relevant imaging depths. Biodistribution studies further demonstrated clear vascular circulation followed by organ–specific accumulation, highlighting the suitability of nJAAZ for time–resolved and deep–tissue imaging applications.

Furthermore, the design features of the nJAAZ platform with the use of ICG and ICG azide and the self-assembly process in physiologically relevant conditions could facilitate its clinical translation. The ability to operate at lower effective dye doses while achieving strong photoacoustic contrast may reduce safety concerns. Moreover, the modular azide–based functionalization strategy increases the versatility of the nJAAZ platform toward identifying disease–specific biomarkers and tissue features^40–42^, paving the way for precision diagnostics in oncology, vascular imaging, cardiovascular diseases, and inflammatory pathologies.

Beyond imaging, the high optical absorption cross–section and tunable surface chemistry of nJAAZ suggest potential extensions into theranostic applications, including photothermal therapy and stimulus–responsive drug delivery^43–48^. Collectively, this study establishes nJAAZ as an adaptable advanced material that bridges supramolecular chemistry, nanomaterials engineering, and biomedical imaging, offering a promising foundation for next–generation photoacoustic contrast agents in translational and clinical settings.

## Materials and Methods

### Materials

The indocyanine green dye (ICG, catalog #91) and the azide-modified indocyanine green dye (ICG-N_3_, catalog #985) are acquired from AAT Bioquest. The dibenzylcyclooctyne-polyethylene glycol 5k (catalog #: PG1-DB-5k), the DBCO-PEG4-biotin (catalog #: PG2-BNDB-5k), and the DBCO-PEG-Folate (catalog #: PG2-DBFA-5k) are purchased from Nanocs. The streptavidin is purchased from Genscript. The Amicon Ultra-0.5 centrifugal filters, the potassium chloride (KCl) salt, the DNA/RNAse free water, the Triton X-100, the dimethyl sulfoxide (DMSO), and the small volume disposable DLS cuvettes are acquired from Sigma-Aldrich. The low melt agarose for electrophoresis was obtained from IBI Scientific. The phosphate buffered saline (PBS) solution at 10X and pH 7.4 was acquired from ThermoScientific and diluted to 1X prior to use with DNA/RNAse free water. The disposable zeta potential measurement cells were purchased from Malvern Panalytical. The India Ink for photoacoustic imaging is purchased from Hardy Diagnostics. The His-tagged human folate receptor 1 (hFOLR1) is acquired from Acro Biosystems. The NTA SPR sensor and the reagents for SPR are acquired from Nicoya Life Science.

### ICG nJAAZ particles formation and purification

The ICG and ICG-N_3_ dye stock solutions are prepared at 5 mg/mL (6 mM) in water and 10 mg/mL (12 mM) in DMSO, respectively, and stored at -20°C. To assemble the JAAZ particles, solutions of ICG dyes at different molar ratios of ICG and ICG-N_3_ (i.e., 1:10, 1:5, 1:3, 2:5, 1:1) are prepared in various KCl concentrations ranging from 1 to 5 mM. The final concentration of total dye (ICG and ICG-N_3_) is fixed at 250 µM (or 1000 µM for the micron size J-aggregates). Unless stated otherwise, all samples are incubated for 18 hours at 60℃ in a thermocycler (BioRad T100) and quickly cooled to 4℃ prior to be filtered using 100 kDa, 0.5 mL centrifugal filters at 4,000g for 10 minutes for three cycles at room temperature, to remove excess of non-incorporated free dye. The filtered nJAAZ particles are then collected by spinning the inverted centrifugal filters at 4,000g for 2 minutes.

### Absorbance measurements

Spin filtered ICG JA particles are diluted to a total dye concentration of 50 µM in 60 µL of water and loaded in a well of a 96-well flat bottom transparent plate. UV-Vis absorbance measurements are recorded from 440-990 nm with a step-size of 2 nm.

### Fluorescence measurements

Spin filtered J-aggregates are diluted to a total dye concentration of 1 µM in 200 µL of water and loaded in a microquartz fluorometer cell. The fluorescence spectra are recorded with a FluoroMax spectrofluorometer (Horiba) after excitation of the nJAAZ at wavelength of 700 nm, 760 nm, and 890 nm.

### ICG concentration and recovery yield quantification

The recovery yield and the total concentration of dye in the samples recovered after filtration is estimated by diluting the JA particles in a 1:5 (v/v) in a solution of 10% Triton X-100 in water and incubated for 10 min at 37°C to disrupt the J-aggregate and releasing the dyes. The estimated amount of free dye in the solution is calculated using a standard curve made of various ratio of ICG and ICG-azide dye to increase accuracy (Figure S14).

### Diameter measurements with dynamic light scattering (DLS)

Filtered J-aggregate formulations are diluted to a total equivalent free dye concentration of 20 µM in 60 µL water and loaded into a low-volume disposable cuvette for DLS. The size measurements are taken using a Zetasizer Nano ZS instrument (Malvern) at a backscattered angle of 173° and a temperature of 25 °C. The measurements are performed at the default viscosity and refractive index of water or PBS depending on the experimental conditions.

### Size, distribution and particles concentration measurements with nanoparticle tracking analysis (NTA)

The size, distribution, and concentration of purified nJAAZ particles diluted in water are measured with NTA using a Particle Metrix Zetaview® instrument using a method previously described^34^.

### Zeta potential measurements

The zeta potential of the JA samples is measured for all formulations at an equivalent free dye concentration of 5 µM in 0.1X PBS at pH 7.4. Briefly, seven hundred µL of the diluted JA particles are loaded into a disposable folded capillary zeta cell (DTS-1070) and zeta potential readings are taken at 25°C using the default viscosity for 0.1X PBS.

### Gel electrophoresis

A 2% (w/v) agarose gel is prepared in 1X Tris acetate EDTA (TAE) buffer in a square gel caster with a 1 mm 9 well comb. A 6 by 6 cm square was cut in the center of 2% gel. A 0.1% (w/v) low-melt agarose gel in 1X TAE buffer is poured into the cut square and allowed to polymerize for 3 hours at room temperature and overnight at 4 °C in a humidity chamber to avoid dehydration. Thirty µL of 350 µM nJAAZ particles are mixed with 10 µL glycerol and loaded into the wells. The gel runs at 120 mV for 15 minutes and is imaged using a blue transilluminator.

### Scanning Electron Microscopy (SEM) and Energy Dispersive X-ray Spectroscopy (EDS)

For SEM, 2 µL of 50 µM nJAAZ particles are deposited on a clean mica surface and air-dried for 24 hours. The samples are then sputter coated with gold and SEM images were collected using a JEOL JSM-IT500HR InTouchScope™ at an accelerating voltage of 10 kV and a working distance of 10 mm. EDS spectra were collected using Octane Elect EDS System at an accelerating voltage of 15 kV and a resolution of 127 eV. ImageJ^49^ (v. 1.54g) was used to determine the size distribution of the particles in the SEM images obtained.

### Fourier Transform Infrared Spectroscopy (FT-IR)

For FT-IR, nJAAZ particles formulated with a 1:5 ratio of ICG-N_3_:ICG in 5 mM KCl and a total dye concentration of 350 µM (after purification) are lyophilized in order to obtain ∼1 mg of powder. The sample is then loaded into a ThermoScientific Nicolet is50 FT-IR spectrometer, and the attenuated total reflectance (ATR) mode is used to analyze the sample. The spectral resolution (4 cm^-1^) and number of scans (32) are imported into the OMNIC software for analysis. The spectral range used to analyze the nJAAZ particle is 400 to 4000 cm⁻¹.

### Mass spectrometry analysis

A SCIEX QTRAP 4500 mass spectrometer is used for the direct infusion analysis of dissolved nJAAZ particles (using Triton X-100 as described in the previous paragraph: “***Quantification of ICG concentration***”) for precise identification and quantification of ICG and ICG-Azide. Sample solutions and standards were prepared at a concentration of 1 µg/mL in methanol and diluted as required to reach the optimal concentration for infusion. The SCIEX QTRAP 4500 is equipped with an electron spray ionization (ESI) spray source in the positive ionization mode. Instrument parameters are carefully optimized for best response, including a source voltage of 4500 volts, a source temperature of 220°C, and gas flows for Gas 1, Gas 2, and curtain gas at 30, 20, and 20 units, respectively. For the infusion process, the prepared samples is loaded into a 1 mL Hamilton syringe or a suitable infusion pump and then steadily infused into the ESI source at a rate of 10 µL/min. The data acquisition phase is conducted, focusing on precursor Q1 scans and product ion (MS^2^) scans across a mass range of m/z = 50-1000. The declustering potential (DP) is 10 volts, entrance potential (EP) is 10 volts, C-Trap exit potential (CXP) is 14 volts, and collision energy (CE) is 40 volts to ensure accurate mass spectrum analysis. The MS/MS spectra are used to verify structures. Finally, the gathered data, encompassing both full scan and MS/MS data, are thoroughly analyzed using the Analyst 1.7 software. Peak area is used for quantification purposes.

### Stability studies of nJAAZ particles

Samples of nJAAZ prepared at 250 µM of equivalent free dye are incubated in 1X PBS (pH 7.4), DMEM supplemented with 100% FBS and serum (100% FBS). Absorbance spectra of the particles are recorded from 440-990 nm at 0, 2, 4, 8, 16, 24 and 48 hours. All particles are incubated at 37°C to mimic physiological temperatures. The change in the intensity of the 895 nm absorbance peak is recorded to determine the stability of the particles. For long term stability, nJAAZ particles formed at 100 µM of equivalent free dye are stored at 4°C in water for up to 185 days. The size and absorbance intensity of the particles at 895 nm is recorded at 0, 8, 20. 38, 53, 88, 108 and 185 days using the procedures mentioned in previous sections.

### Functionalization of nJAAZ particles

The nJAAZ particles prepared at different ratios of ICG dyes are functionalized with various functional groups following a similar protocol described below.

The azide groups on nJAAZ particles following filtration are functionalized with DBCO-polyethylene glycol (PEG) to increase their stability in physiological conditions. For DBCO–PEG conjugation, the number of functional azide groups on the nJAAZ particles is determined by estimating the free dye concentration. For example, a 1:1 ICG–azide:ICG formulation at an estimated free dye concentration of 300 µM corresponds to approximately 150 µM of available azide groups. Ten percent of these available azide groups (in µM) are subsequently modified with DBCO–PEG5k using a stock solution prepared at 10 mg/mL in DMSO. The reaction is left at room temperature for 3 hours. The excess of DBCO-PEG5k is removed through filtration using 100 kDa centrifugal filters, as mentioned above.

To prove the versatility of functionalization of the nJAAZ particles, the particles are functionalized with streptavidin using the same functionalization scheme established in our previous work^33^. nJAAZ particles with 1:3 ICG-azide:ICG ratio are labelled with DBCO-PEG4k-biotin at 10 times the concentration of available azide groups. The reaction occurred for 18 hours at 4 °C. Excess DBCO-biotin was removed through filtration using 30 kDa centrifugal filters. Following removal of excess biotin, the amount of estimated free dye is calculated as mentioned in the previous section. The relative concentration of available biotin groups is equivalent to the concentration of ICG-azide dye, calculated from the estimated free dye concentration. Following biotin addition, the particles are functionalized with streptavidin at 1.5-fold the concentration of available biotin groups, at room temperature for 30 minutes. Excess streptavidin is removed by filtration using 0.5 mL, 100k Da centrifugal filters. The particles are filtered 4 times at 4000*g* for 10 minutes.

To create a platform for the attachment of proteins, nJAAZ particles were functionalized with DBCO-PEG3k-NTA. nJAAZ particles with different ratios of ICG-N_3_:ICG (1:10, 1:5, 1: 3, 2:5 and 1:1) were conjugated with 5X the concentration of available azide groups for each ratio. The reactants were conjugated for 18 hours at 4 °C. The stock of DBCO-PEG3k-NTA was at 10 mg/mL in DMSO. Unreacted molecules are removed through filtration using 30kDa filters.

To create targeted particles, DBCO-PEG5k-Folate is functionalized onto nJAAZ particles with 1:3 or 2:5 ICG-azide:ICG ratios. A 20 mg/mL DBCO-PEG5k-Folate stock is diluted to functionalize 10% of the estimated azide groups present after filtration. For example: The concentration of azide groups in 1:3 ICG-azide:ICG particles at 300 µM estimated free dye concentration is 90 µM. 10% (in µM) of the 90 µM available azide groups is modified with the folate group. Complete functionalization of all available azide groups with a DBCO-PEG5k-folate increased the size of the particles by ∼30%. Hence, 10% is chosen as the optimum concentration for functionalization. Excess and unreacted DBCO-Folate is removed through filtration using 30 kDa filters. To ensure complete removal of the targeting moiety, the samples are filtered for 10 minutes at room temperature for 4 cycles of 4 minutes each.

### Surface plasmon resonance (SPR) studies

The nJAAZ-FA binding curves are obtained with an Open Surface Plasmon Resonance (Nicoya Lifesciences) using the NTA sensor chips modified with the His-tagged human folate receptor 1 (FOLR-1). Prior to running the SPR measurements, the NTA surface is regenerated with two injections of 100 mM EDTA injection (150 µL/min) and activated with one injection of 40 mM NiCl_2_ at 20 µL/min. The FOLR1 is then injected at 25 µg/mL prior to injecting various concentrations of nJAAZ. The surface is regenerated with 100 mM of EDTA.

### Breast cancer cell culture, targeting, and immunofluorescent staining

Estrogen Receptor-positive/HER2-negative ER+/HER2-breast cell line T47D was obtained from the American Type Culture Collection (ATCC). As per manufacturer guidelines, cells are cultivated in RPMI-1640 medium supplemented with 10% fetal bovine serum (ATCC, Manassas, VA, USA) and human recombinant insulin in zinc solution (Thermo Fisher Scientific, Waltham, MA USA) in a humidified atmosphere with 5% CO_2_ at 37°C. Based on cell growth rate, cells are sub-cultured using trypsin/EDTA (ATCC, Manassas, VA, USA) at ratios ranging from 1:3 to 1:6. For imaging, a total of 40,000 cells were seeded per well into an 8-well chambered coverslip (Ibidi, Fitchburg, Wisconsin, USA) and allowed to grow for 24 hours. Wells are then washed twice with ice cold PBS before fixing and permeabilizing with methanol for 15 minutes on ice and blocked with 5% bovine serum albumin (BSA) in PBS supplemented with 0.1% Tween-20 for 1 hour. Samples are incubated overnight at 4°C with nJAAZ particles diluted at 1:400 in PBS supplemented with 1% BSA. Finally, cells are stained with DAPI to visualize the nucleus and imaged on a CrestOptics X-Light Series X-Light V2 Spinning Disk Confocal microscope (Nikon Instruments Inc., Melville, NY, USA). The Cy5 signal was captured at 100 ms exposure for all samples and DAPI was captured at an exposure of 1 ms, and images were processed with 2D deconvolution before exporting as TIFF files.

### Characterizing the Photoacoustic properties of the nJAAZ in vitro

The PA properties of the different formulations of nJAAZ were evaluated as previously described using our homemade PA setup^33^. Briefly, the nJAAZ samples embedded in 2% agarose are loaded in a 96-well plate and placed in a plastic reservoir filled with deionized water to fully cover the plate. We are using a 35 MHz (PI35, Olympus, Massachusetts, USA) or a 5 MHz (V326, Olympus, Massachusetts, USA) focused transducer submerged in water and at a distance of 12 mm or 45 mm from each sample loaded in the plate. Sample positioning and PA signal optimization are achieved by scanning each sample with a fine x–y translation of the stage located above the samples and where the transducer and the pulsed laser remain fixed. Samples are excited obliquely using a wavelength–tunable pulsed laser (690–950 nm; Phocus Mobile, Opotek) at 895 nm. The transducer signal is amplified (20/40 dB; HVA–200M, Femto) and recorded using a lock–in amplifier (Zurich Instruments) synchronized to each laser pulse via a photodiode. For each measurement, 10 s of data (100 waveforms) are collected per sample or per wavelength for spectral measurements. The absolute amplitude of each waveform is calculated, averaged, and normalized to the signal from India Ink (1:100 ratio) or whole sheep blood containing anticoagulant. The nJAAZ samples were tested at 5, 12, and 25 µM. For the “M” shape PA imaging, HeLa cell cultured at 37 °C and 5% CO2 in Eagle’s minimum essential medium (EMEM) complemented with 10% Fetal Bovine Serum (FBS) and 1% Penicillin are seeded at a concentration of 1.0 × 10^5^ cells/ml on a glass slide where a 3D printed “M” shape has been placed to prevent cell adhesion. After removing the 3D printed “M” shape, the cells are incubated with 10 µM nJAAZ or targeted nJAAZ-FA for 20 min and then washed to remove unbound nJAAZ. The surface of the slides are then raster scanned via stage translation with 100 µm steps with the same set previously described above. The sample was then raster scanned through movement of the stage at 100 µm steps. The images are created by plotting the absolute signal magnitude in 2D space and further smoothen via a Matlab image gaussian filter.

### In vivo studies and PA image acquisition

Male 4-6-week-old athymic Swiss nu/nu nude mice (Jackson Laboratories, Bar Harbor, ME) are used for these studies. Mice are housed (5 mice per cage) in a room maintained at constant temperature and humidity under a 12-hour light and dark cycle. Mice are fed a regular chow diet with access to water *ad libitum*. All experimental protocols are reviewed and approved by the Institutional Animal Care and Use Committee (IACUC) at UTHealth (IACUC #AWC-22-0098). Mice are anesthetized with isoflurane (1-2.5%) mixed with oxygen (1-2 L/min) and immersed in the PhotoSound TriTom® imaging device chamber filled with deionized and degassed water and maintained at a temperature of 36°C. Mice are suspended vertically in a custom-jig that allowed inhalation of anesthetic while the head stayed above water. Various concentrations of nJAAZ particles are administered intravenously immediately before immersion in the water tank and continuous imaging is performed for up to 60 minutes. The TriTom system includes an Ekspla PhotoSonus M-20 laser (3-5 ns pulse duration, and 20 Hz pulse repetition rate; Ekspla, Vilnius, Lithuania) for optical signal generation via four optical fiber terminals mounted vertically around the water tank at 45 and 90 degrees with respect to a 6 MHz central frequency arc transducer array with 96 channels for acoustic signal acquisition. The vertical suspended mouse was rotated at 360 degrees by a stepper motor attached to the nose cone that delivers isoflurane. The spatial resolution of TriTom is 173 ± 17 μm in the transverse plane and 640 ± 120 μm in the longitudinal axis with an absorption coefficient sensitivity detection limit of 0.258 cm-1, and an imaging speed of 38 s. 3D images and 2D cross sectional projections obtained during imaging at 895 nm, the optical absorption wavelength of nJAAZs, are reconstructed using a standard modified back-projection algorithm (TriTom reconstruction software V3.0.3) and additional processing steps are performed in MATLAB V2021b as described previously^50^.

### In vivo Contrast-to-Noise ratio and imaging depth

Multiwavelength imaging is performed on multiple mice at three timepoints (T0, T30, and T60 minutes) and for various nJAAZ concentrations. As a reference, pre-injection images are also acquired. The acquired photoacoustic data is reconstructed into image volumes using the Tritom Reconstruction Software (TRS). Reconstructed volumes are output into VTK and MAT files for further analysis. The VTK volumes are loaded into 3D slicer where coronal maximum intensity projections of the mouse trunk are generated for different time points. The volumes are then loaded into MATLAB where 0.5 mm cross-sectional maximum intensity projections were generated at 2 different axial positions to highlight anatomical features. Linear unmixing is then performed with the TRS to unmix signals from oxyhemoglobin, deoxyhemoglobin, and nJAAZ using the multi-wavelength acquisitions. Spectral unmixing for oxyhemoglobin (red), deoxyhemoglobin (blue), and nJAAZ (green) is done for the reconstructed volume using multiple wavelengths: 700, 730, 760, 800, 850, 875, and 895 nm. For referencing anatomical structures, the unmixed volume is overlaid with a mixed 895 nm scan. Two additional 0.5 mm maximum intensity projections are generated to highlight organs and the iliac arteries. For quantitative analysis, Contrast-to-Noise ratio (CNR) and imaging depth are found for the iliac artery at the 60-minute timepoint post injection of the nJAAZ particles at a concentration of 150 µM. The PA signal intensity of the iliac artery and a reference background are averaged respectively across three axial slices containing the iliac arteries. The CNR is calculated using the following equation:

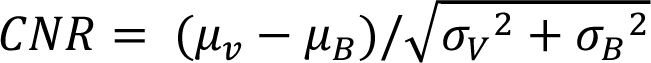

Where µ_v_, µ_B_, σ_v_, and σ_B_ are the mean and standard deviation of the vessel and background respectively. Imaging depth of the artery is reported by measuring the shortest distance from the vessel to the skin line of mouse.

### mJAAZ and nJAAZ in vitro photoacoustic signal intensity comparison

Two samples are prepared with molar ratios of ICG-N_3_:ICG of 1:10 and 1:1 in 20 mM KCl, corresponding to mJAAZ (micron-sized JAAZ, 1120 nm) and nJAAZ (148 nm), respectively. Photoacoustic data for these samples are collected using the Fujifilm Vevo F2. For each sample type, measurements are performed at two concentrations (15 µM and 50 µM estimated total dye concentration), with two independent replicates per concentration. For each replicate, a fresh sample is prepared and loaded into new polyurethane (PU) tubing to avoid reuse. Samples are stabilized inside a Vevo Contrast Agent Phantom filled with deionized (DI) water. A 29 MHz transducer (center frequency: 20 MHz; bandwidth: 15–29 MHz) is used for all measurements, and the transducer’s fiber jacket cavity is filled with degassed clear ultrasound gel. Using Spectro mode, the absorption profile of each contrast agent is first measured to determine the peak absorbance wavelength for acquisition, which is 865 nm for both samples. Photoacoustic signal intensity is then acquired for 20 seconds (100 frames) and averaged along the length of the tubing. Raw data analysis and quantification are conducted using Vevo software. Signal intensity is measured within a consistent region of interest (ROI) of 2.57 mm² across frames.

## Acknowledgement

G.T. acknowledge the Meyerhoff Scholar’s program. Funding for this research was provided via two Virginia Innovation Partnership Corporation (VIPC), Commonwealth Commercialization fund grant for Higher education award #CCF23-0092-HE (tier 1) and #CCF26-0003-HE2 (tier 2) to P.V.C. and R.V., and a seed grant from the Office of Research, Innovation, and Economic Impact (ORIEI) at George Mason University (award #G00002563) to J.L.M., M.P., P.V.C., and R.V.

## Author contribution

Conceptualization, P.V.C., R.V.; Methodology, S.S., L.S.C., E.R., M.G., M.P., S. K., P.V.C, and R.V.; Investigation, S.S., L.S.C., N.S., M.H., G.G., D.J.L., I.E.H., B.C., I.F., G.T., D.V.S., E.L.G., E.R., M.G., and R.V.; funding acquisition, J.L.M., M.P., P.V.C, and R.V.; S.S. and R.V. wrote the first draft of the manuscript. All authors contributed to editing the manuscript. All authors have given their approval to the final version of the manuscript.

