## Supplementary Information for "Direct Synthesis of Targeted Nanosized ICG J-aggregate for Photoacoustic Imaging"

**Table S1**: Polydispersity Index (PDI) and size (nm) of the different molar ratio ICG-$N_3$:ICG samples tested.

| Molar ratio ICG-$N_3$:ICG | Molarity of KCl [mM] | | | | |
|---|---|---|---|---|---|
| | 1 | 2 | 3 | 4 | 5 |
| 1:10 | 0.462 ± 0.066 | 0.423 ± 0.033 | 0.533 ± 0.077 | 0.533 ± 0.077 | 0.581 ± 0.095 |
| 1:5 | 0.413 ± 0.093 | 0.478 ± 0.098 | 0.727 ± 0.075 | 0.721 ± 0.082 | 0.691 ± 0.194 |
| 1:3 | 0.168 ± 0.022 | 0.163 ± 0.02 | 0.175 ± 0.016 | 0.202 ± 0.012 | 0.210 ± 0.024 |
| 2:5 | 0.124 ± 0.023 | 0.143 ± 0.023 | 0.151 ± 0.024 | 0.158 ± 0.013 | 0.143 ± 0.019 |
| 1:1 | 0.079 ± 0.011 | 0.087 ± 0.026 | 0.116 ± 0.019 | 0.127 ± 0.013 | 0.109 ± 0.016 |

n=3 distinct samples; data is represented as mean±SD.

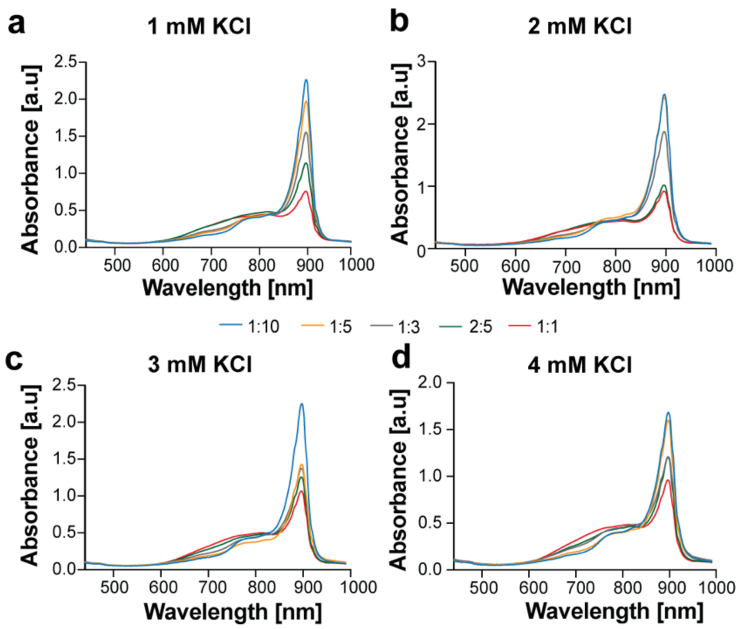


**Figure S1**: UV-Vis-NIR spectrum of different molar ratios of ICG-azide:ICG namely 1:10, 1:5, 1:3, 2:5 and 1:1 in (a) 1 mM KCl, (b) 2 mM KCl, (c) 3 mM KCl and (d) 4 mM KCl. All absorbance readings were taken at a volume of 60 µL (n=3 distinct replicates)

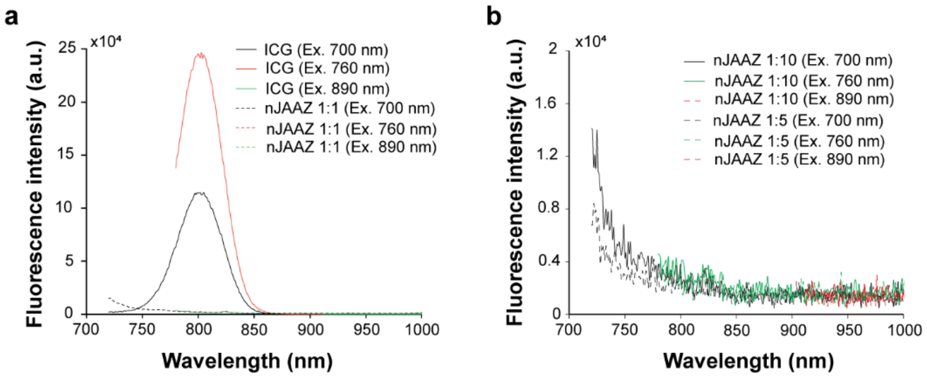


**Figure S2**: Representative fluorescence spectroscopy curves of ICG and nJAAZ at 1:1 ratio (a) or at 1:10 ratio (b) and at 1 μM ICG or total ICG dye concentration.

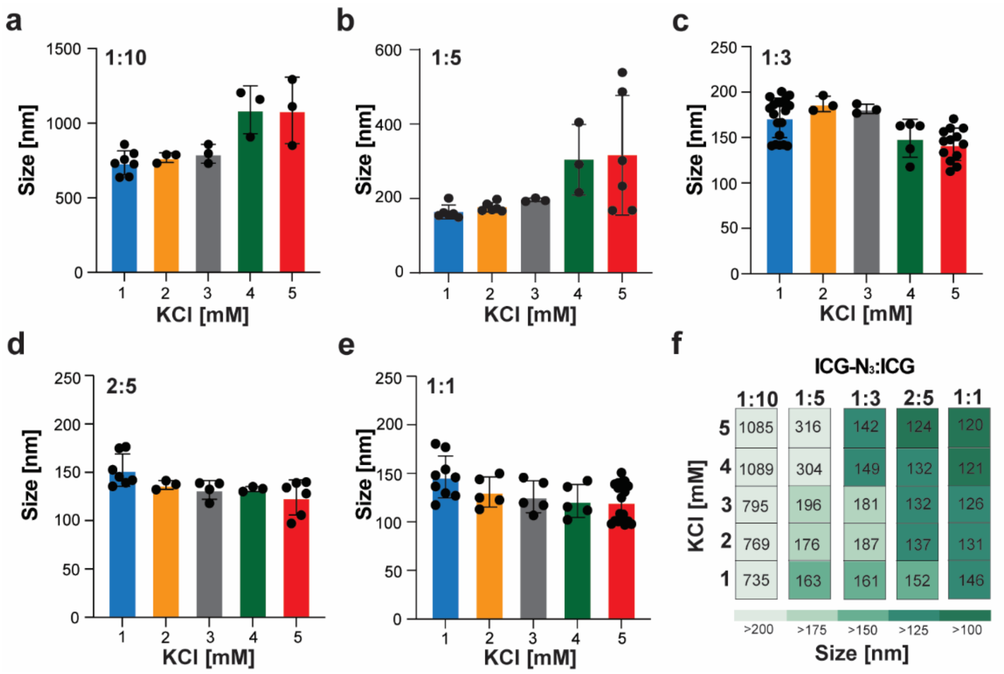


**Figure S3**: DLS measurements of the Hydrodynamic diameters of J-aggregates synthesized with (a) 1:10, (b) 1:5, (c) 1:3, (d) 2:5 and (e) 1:1 ICG-azide:ICG molar ratios with different KCl molarities namely 1, 2, 3, 4 and 5 mM KCl. The total concentration of dye in all solutions was 350 $\mu$M. (n>3 distinct replicates, with error bars representing the standard deviation) (f) A schematic summary of the different sized J-aggregates synthesized with varying salt and ICG-$N_3$:ICG molar ratios.

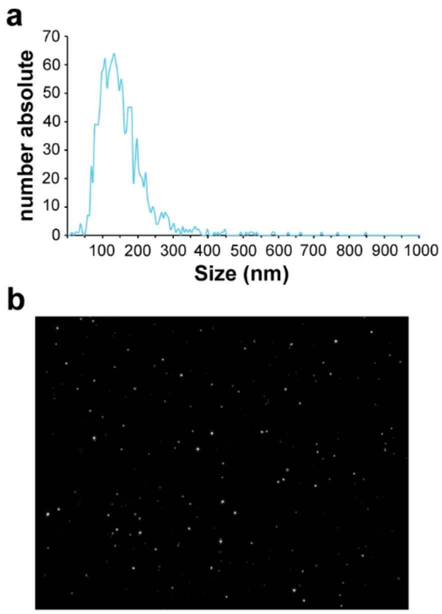


**Figure S4**: Representative Nanoparticle tracking analysis (NTA) measurement results using the ZetaView® system with nJAAZ formed at a 1:1 ratio and 5 mM KCl. (a) Number weighted size distribution. (b) Representative single frame captured from the video used to measure the size of the nJAAZ.

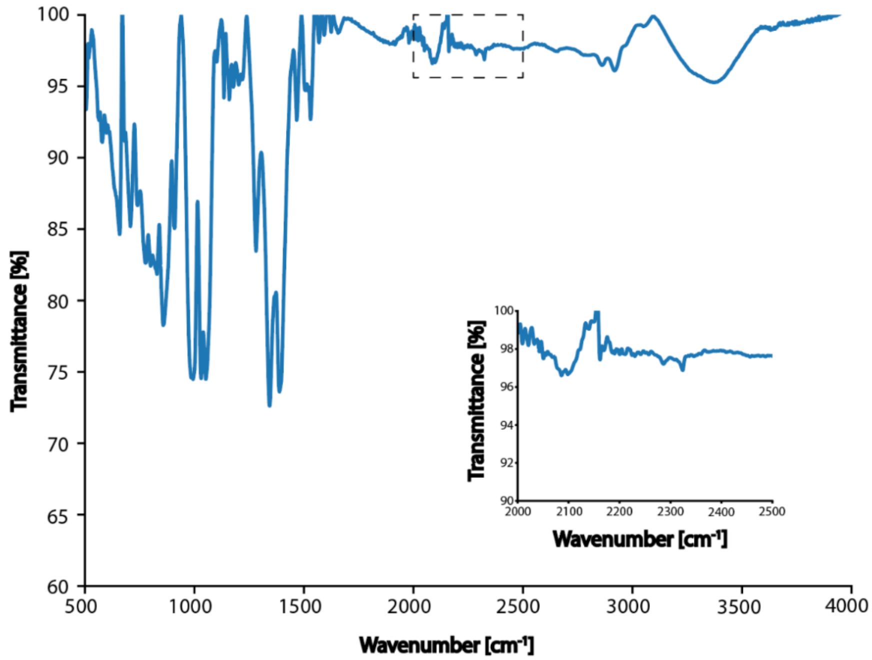


**Figure S5**: FT-IR spectra of J-aggregates synthesized with a 1:5 ICG-$N_3$:ICG molar ratio using 5 mM KCl. The zoomed in figure displays a peak at 2100 $cm^{-1}$ to show the presence of azide.

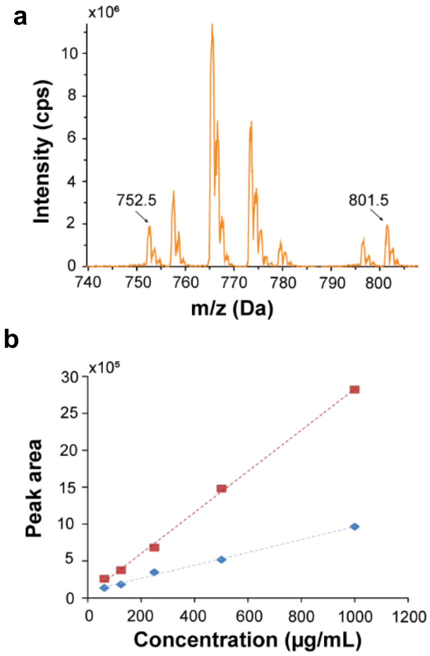


**Figure S6**: Mass spectrometry characterization of the J-aggregates formed with 1:1 ICG-$N_3$:ICG ratio (5 mM KCl). (a) Representative total ion Chromatogram showing the ICG peak (m/z=752.5) and the ICG Azide peak (m/z=801.5). (b) Representative calibration curve showing the instrument response (peak area) versus the concentration of both ICG (blue) as well as ICG-$N_3$ (red).

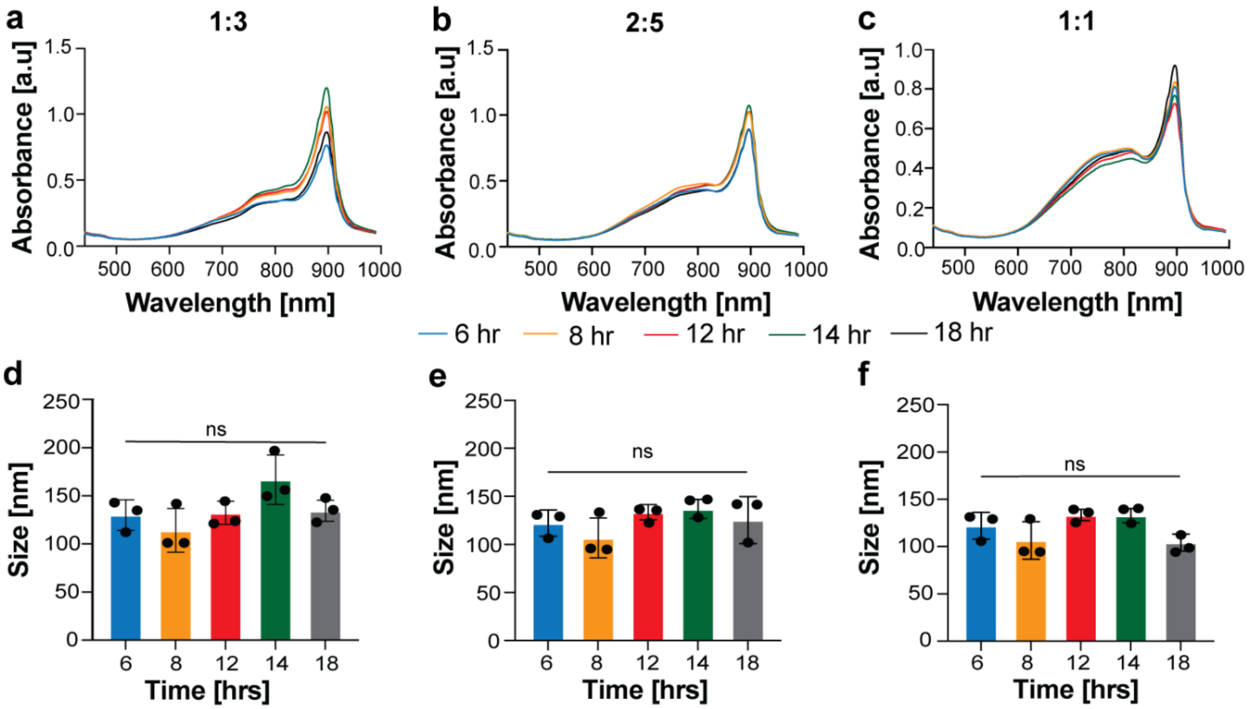


**Figure S7**: UV-Vis-NIR spectra of (a) 1:3, (b) 2:5, and (c) 1:1 ICG-$N_3$:ICG molar ratio samples formed after 6, 8, 12 and 18 hours of incubation in 5 mM KCl. (d-f) Average size of 1:3, 2:5 and 1:1 molar ratio sample after 18 hours of incubation. (n=3 distinct replicates)

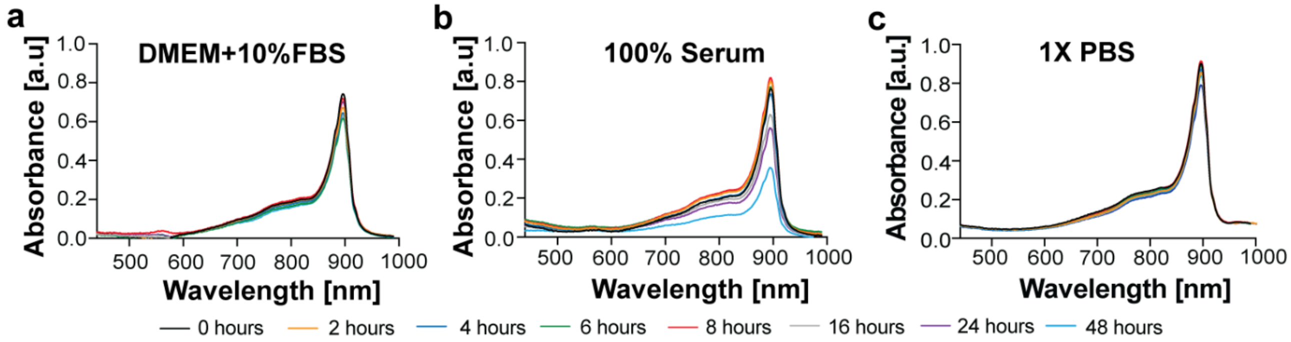


**Figure S8**: UV-Vis-NIR spectrum of nanosized J-aggregates incubated in (a) DMEM+10%FBS, (b) 100% FBS and (c) 1XPBS for 48 hours at 37 ℃. All absorbance readings were taken at an equivalent dye concentration of 15 $\mu$M and 60 µL volume. (n=3 distinct replicates)

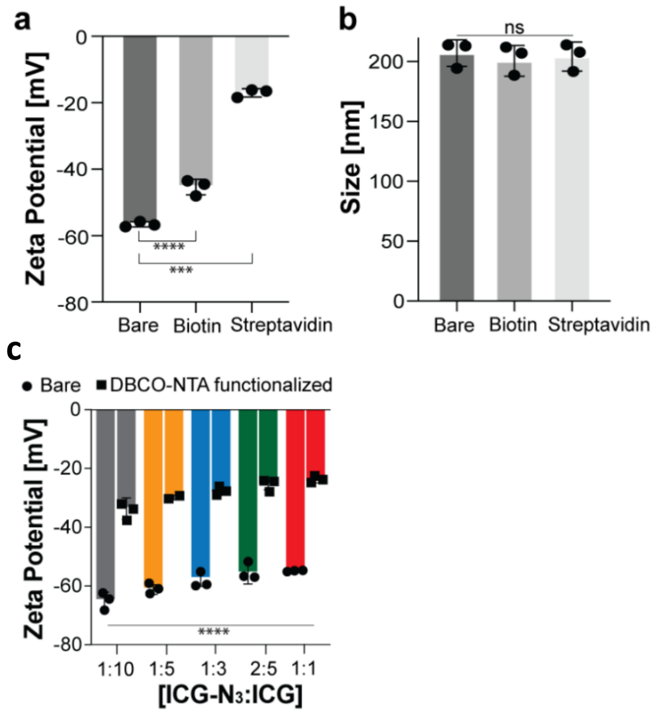


**Figure S9**: (a) Zeta potential of 1:3 ICG-$N_3$:ICG molar ratio nJAAZ particles modified with biotin followed by streptavidin. (b) Hydrodynamic diameter of nJAAZ particles after functionalization with biotin and streptavidin. (c) Zeta potential measurements of different molar ratio nJAAZ particles before *(circle)* and after functionalization *(square)* with DBCO-NTA.

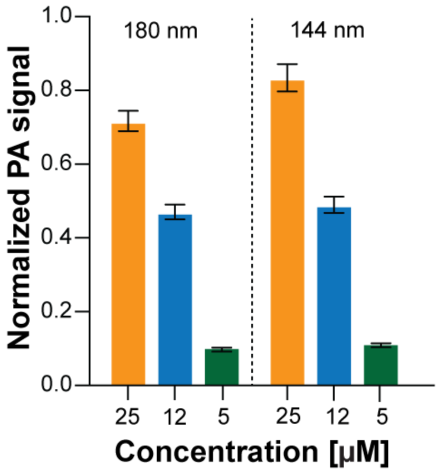


**Figure S10**: Photoacoustic signal amplitude of nJAAZ particles of two different hydrodynamic sizes which are 180 nm and 144 nm at different equivalent concentrations of free dye. The data is normalized with respect to 1:100 ratio of India Ink used as a control.

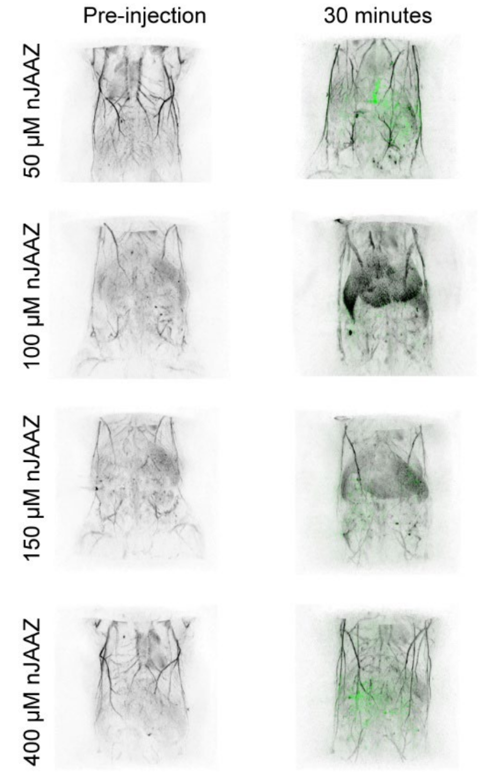


**Figure S11**: Coronal slab maximum intensity projections of mouse trunk pre-injection and 30 minutes post-injection for nJAAZ particle concentrations 50 µM, 100 µM, 150µM, and 400 µM. For 30 minutes post-injection, nJAAZ signal was spectrally unmixed and shown in green.

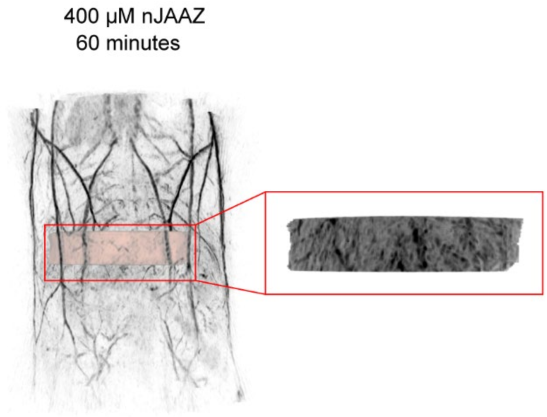


**Figure S12**: Coronal slab maximum intensity projection of mouse trunk 60 minutes post-injection with nJAAZ particle concentration of 400μM. A section of the liver was segmented with an adjusted dynamic range and is highlighted with a red outline.

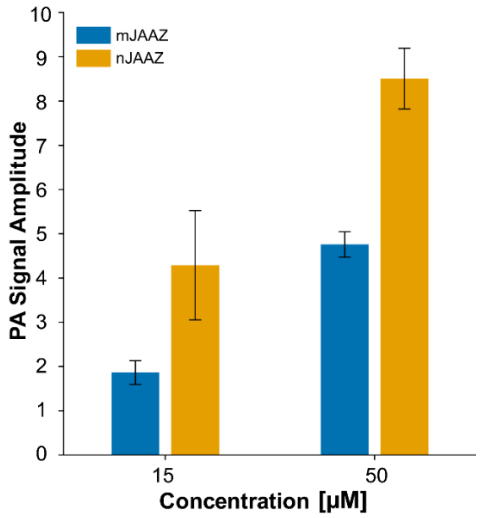


**Figure S13**: Photoacoustic signal intensity of nJAAZ and mJAAZ samples at two different equivalent concentrations of free dye (15 and 50 $\mu$M). The hydrodynamic size of each sample used is 148 nm and 1120 nm, respectively for nJAAZ and mJAAZ. For total dye concentrations listed, nJAAZ shows a higher signal amplitude compared to mJAAZ although standard error bars indicate lower signal variations for mJAAZ (n=2).

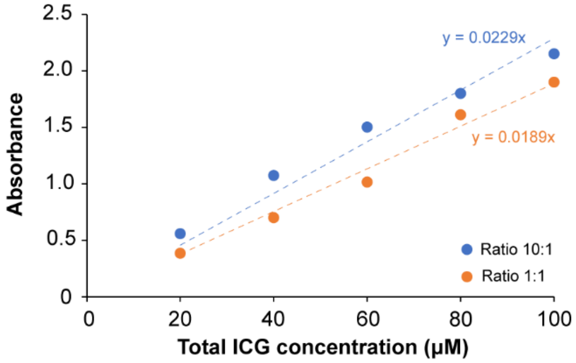


**Figure S14**: Representative standard curve of free ICG and ICG azide dye at two different ratios (1:10 [blue] and 1:1 ICG-$N_3$:ICG [orange] ratios).
